# 3D Histology Validates 2D Histology for Axon Radius Distributions and Conduction Velocities

**DOI:** 10.64898/2026.03.25.714137

**Authors:** Laurin Mordhorst, Nikolaus Weiskopf, Markus Morawski, Siawoosh Mohammadi

## Abstract

Axons are the brain’s wiring, organized into bundles that connect nearby and distant regions. Axon caliber determines signal conduction velocity and varies both within and across bundles, reflecting the brain’s diverse functional demands. Much of what we know about this organization derives from 2D histology, assuming cylindrical axons whose calibers are described by their radius. Yet, recent 3D histology reveals that the radius varies along an individual axon—with implications for both characterizing axon caliber and potentially conduction velocity predictions. We show in 450,000 3D rat axon reconstructions that—despite this individual variation—axon bundles possess stable radius distributions at the ensemble level, which 2D cross-sections faithfully represent. This representativeness extends to conduction velocity predictions, as along-axon variation has only modest impact. In particular, large axons exhibit especially stable conduction, emphasizing their key role in time-critical signaling. With 2D sampling validated, we leverage 46 million human corpus callosum axons from 2D histology to determine sample size requirements across neuroscience applications. Our findings reinforce decades of 2D histology-based research on axon organization and its functional implications, while guiding future study design.

## 1 Introduction

The brain’s white matter connects distant regions via tracts—bundles of aligned myelinated axons. The caliber of these axons varies both within and across tracts, and is a key determinant of conduction velocity: thicker axons conduct faster, enabling rapid long-range communication (Ruthig et al., 2025), while thinner axons are more numerous and metabolically efficient, serving less time-critical communication (Perge et al., 2012). Beyond basic neuroscience, axon caliber holds promise as biomarkers for neurological disorders (Evangelou et al., 2001; Stassart et al., 2018) and is in principle accessible non-invasively via magnetic resonance imaging (MRI) (Alexander et al., 2010; Assaf et al., 2008; Mordhorst et al., 2025a; Veraart et al., 2020).

Much of our understanding of axon organization derives from 2D histological cross-sections, interpreted under the cylinder model—the assumption that axons are uniform tubes characterized by their radius (Aboitiz et al., 1992; Caminiti et al., 2009; Graf von Keyserlingk and Schramm, 1984; Liewald et al., 2014; Wegiel et al., 2018). This approach has yielded a rich body of findings. Axon radius distributions are right-skewed, with a large bulk of many thin axons and a sparse tail of few thick, fast-conducting axons—a shape often captured by parametric distributions such as the gamma or generalized extreme value distribution (Assaf et al., 2008; Lee et al., 2019; Oliveira et al., 2023; Sepehrband et al., 2016). While the bulk of the distribution is similar across species, the length of the distribution tail varies, with larger brains extending the tail to include giant axons, partially accounting for longer conduction distances (Caminiti et al., 2009; Olivares et al., 2001). Within brains, tracts differ systematically: motor-related pathways contain larger axons than association tracts (Liewald et al., 2014; Tomasi et al., 2012), and long-range axons are thicker than short-range axons (Ruthig et al., 2025). Additionally, 2D histology-derived distributions have shown consistency with functional and biophysical measurements grounded in the cylinder model, including conduction velocity measurements from invasive electrophysiology (Salami et al., 2003; Skoven et al., 2025) and human electroencephalography (Oliveira et al., 2023; Ulusoy et al., 2004), as well as axon radius measurements with diffusion MRI (Mordhorst et al., 2025a; Veraart et al., 2020).

Yet, the consistency of these findings may be surprising given known limitations of 2D histology. First, most studies use microscopy sections much narrower than the tracts they sample, with sample sizes potentially insufficient to characterize distribution tails or validate parametric radius distribution models. Second, 3D axon reconstructions from serial-section electron microscopy have revealed that individual axons deviate from the cylindrical model, exhibiting radius variation and undulations along their trajectory (Abdollahzadeh et al., 2019, 2021; Andersson et al., 2020; Lee et al., 2019; Shapson-Coe et al., 2024; Tian et al., 2025). Though there are hints that axon radius distributions remain stable along tracts based on few (∼50) large axons (Andersson et al., 2020), this remains to be established quantitatively at scale. Beyond histological sampling, deviations from cylindrical geometry affect biophysical modeling with diffusion MRI (Andersson et al., 2020, 2022; Lee et al., 2019, 2020, 2024; Nilsson et al., 2012; Winther et al., 2024) and conduction velocity predictions (Kolaric et al., 2013). For conduction velocity, it is of particular interest whether axons of different sizes are differentially affected, given their distinct functional roles (Perge et al., 2012; Wang et al., 2008). Under what conditions, then, do 2D histology-derived axon radius distributions yield meaningful functional and biophysical predictions?

Here, we study along-axon radius variation and its implications, the robustness of axon radius distributions to histological sampling (2D vs 3D, sample size), and their parametric description. Leveraging the largest available dataset of 3D rat axon reconstructions (450,000 corpus callosum and cingulum axons), we first show that radius variation along axons has modest impact on conduction velocity, with large axons enabling particularly stable conduction. We then mimic 2D histological sampling and find that 2D-derived radius distributions faithfully represent 3D bundle properties despite radius variation along individual axons. With 2D histological sampling validated, we turn to the largest available 2D human dataset (46 million corpus callosum axons) to provide sample size guidance: even small samples (∼10^3^ axons) reasonably capture the distribution bulk, whereas larger samples (∼10^5^ axons) are needed to capture the long distribution tail in humans. Finally, we show that this long tail challenges parametric modeling of axon radius distributions.

## 2 Results

We extracted axon radii from two complementary deep white matter datasets (see Fig. 1): 3D electron microscopy of rat corpus callosum and cingulum (∼450,000 reconstructed axons across 19 ROIs, see Fig. 1a), and 2D light microscopy of human corpus callosum (∼46 million axons across 35 ROIs, see Fig. 1b). The 3D data enables analysis along individual axon trajectories; the 2D data provides extensive sampling across distinct axons.

**Figure 1.**
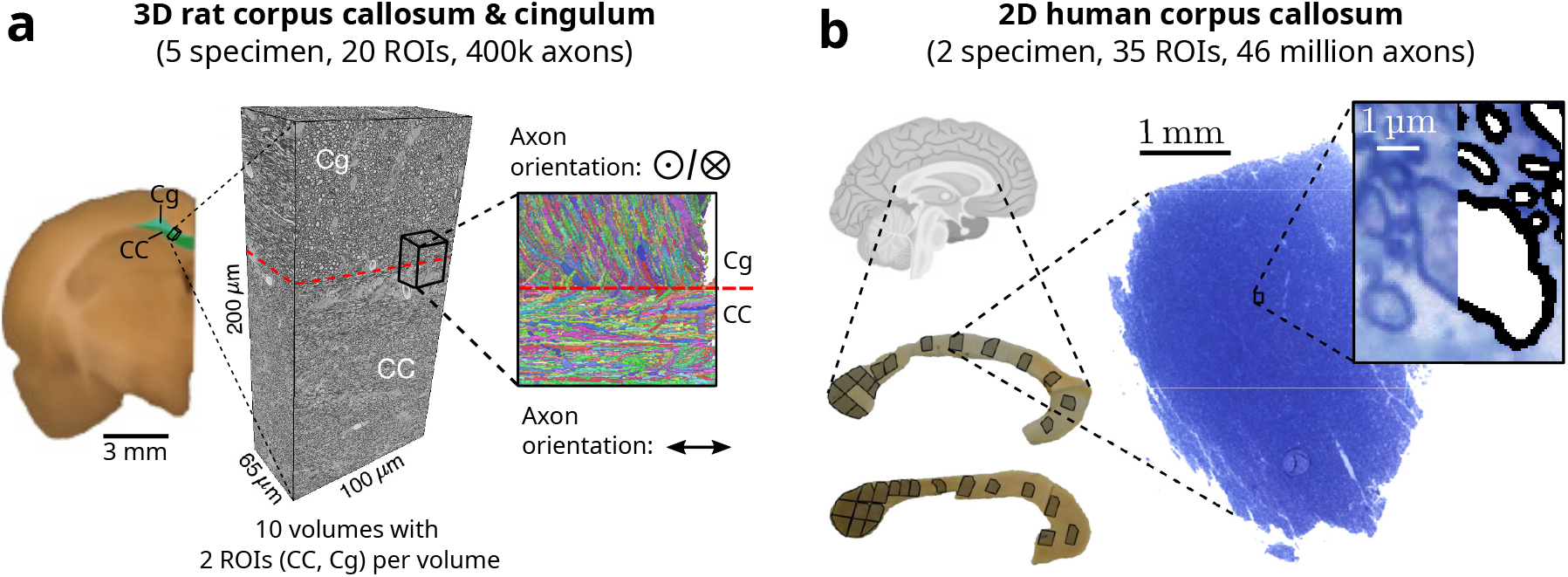
Dataset overview. **(a)** 3D electron microscopy dataset from rat white matter. Left: coronal view of rat brain hemisphere showing the location of the electron microscopy volume spanning corpus callosum (CC) and cingulum (Cg). Center: representative electron microscopy volume. Right: 3D axon segmentations for a subvolume with main axon orientation indicated; each color represents a distinct axon. Dataset comprises 19 ROIs distributed over 10 volumes from 5 rats (2 sham-operated controls, 3 after traumatic brain injury), totaling approximately 450,000 reconstructed axons. **(b)** 2D light microscopy dataset from the human corpus callosum. Left: anatomical schematic showing ROI locations across two post-mortem samples. Right: representative light microscopy image with inset illustrating axon segmentation. Dataset comprises 35 ROIs totaling 46 million segmented axons. Rat brain schematic and electron microscopy volume in (a) were adapted from Abdollahzadeh et al. (2025), licensed under CC BY 4.0.

### Along-axon radius variation has modest impact on conduction velocity

To characterize radius variation along axons, we extracted radius profiles from the 3D axon reconstructions in rats. Fig. 2a shows 3D renderings of three representative axons with low, medium, and high along-axon radius variation; Fig. 2b shows the corresponding radius along the axonal trajectories. The radius changes gradually along axons, with the thickest axon (green) showing less variation than thinner ones (orange, purple).

**Figure 2.**
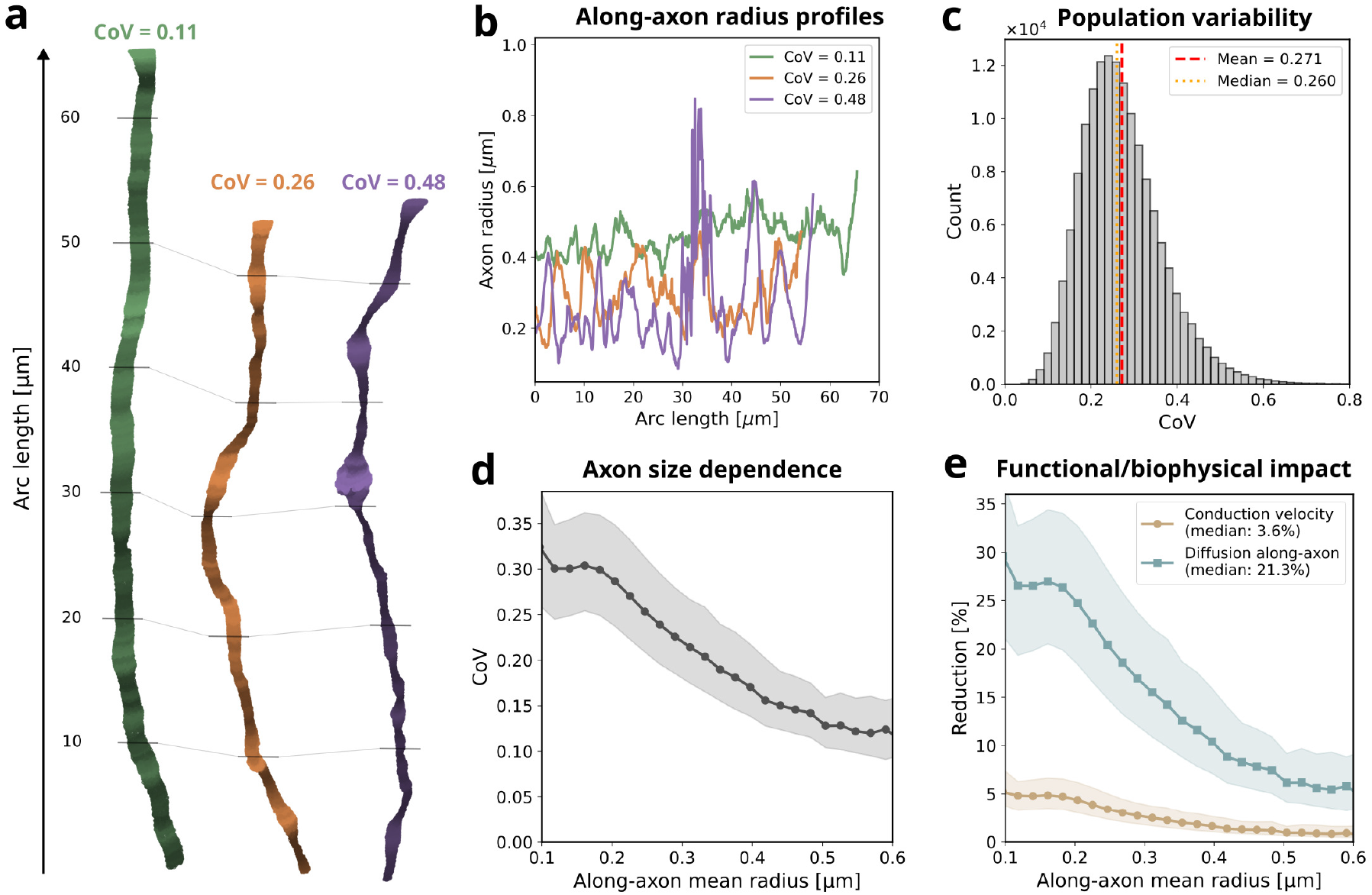
Along-axon radius variation and its impact. **(a)** 3D renderings of three representative rat corpus callosum axons with different levels of along-axon radius variation. Variation is quantified by the coefficient of variation (CoV; see Eq. (1)), annotated above each axon. Brightness indicates local radius (dark = low, bright = high). **(b)** Radius profiles along arc length for the axons in (a), with radius measured perpendicular to the centerline. **(c)** Distribution of CoV across all axons; dashed lines indicate mean and median. **(d)** CoV as a function of along-axon mean radius. Axons are binned by along-axon mean radius; line shows median, shaded region shows interquartile range (IQR). **(e)** Estimated reduction of conduction velocity (see Eq. (4)) and along-axon diffusion (see Eq. (7)) due to radius variation; 0% corresponds to ideal cylindrical axons without radius variation (CoV = 0). Data are displayed as in (d). The legend provides the median reduction across all axons. Statistic computation in (c–e) was restricted to axons exceeding 20 µm arc length (*n* ≈ 150,000) to robustly quantify along-axon CoV; bins in (d–e) were constrained to a minimum of 50 axons.

Moving from representative axons to population level, Fig. 2c shows the histogram of alongaxon radius variation as coefficient of variation (CoV), i.e., the standard deviation of the radius relative to the along-axon mean (see Eq. (1)). The CoV has a median of 0.27, but can reach up to 0.6 for some axons. Interestingly, CoV shows a clear dependency on the mean along-axon radius (see Fig. 2d), decreasing for thicker axons. Fig. S1 reveals the mechanism behind this: the absolute radius variation saturates for thick axons.

To assess the functional and biophysical impact of radius variation (see Fig. 2e), we estimated the CoV-related reduction of conduction velocity and along-axon diffusion—accessible in vivo with non-invasive diffusion MRI—using Eqs. (4) and (7). Conduction velocity is much less affected than along-axon diffusion, with median reductions of ∼4 % and ∼21 % respectively.

### 2D cross-sections capture axon radius distributions

While individual axon radii vary along their trajectory (see Fig. 2), the axon ensemble’s radius distribution along the bundle may nonetheless remain stable. We evaluated this hypothesis by tracing local axon radius distributions along bundles, using cross-sections mimicking 2D histological sampling, and compared them to the full 3D distribution, which pools radii along and across all axons (see Fig. 3a).

**Figure 3.**
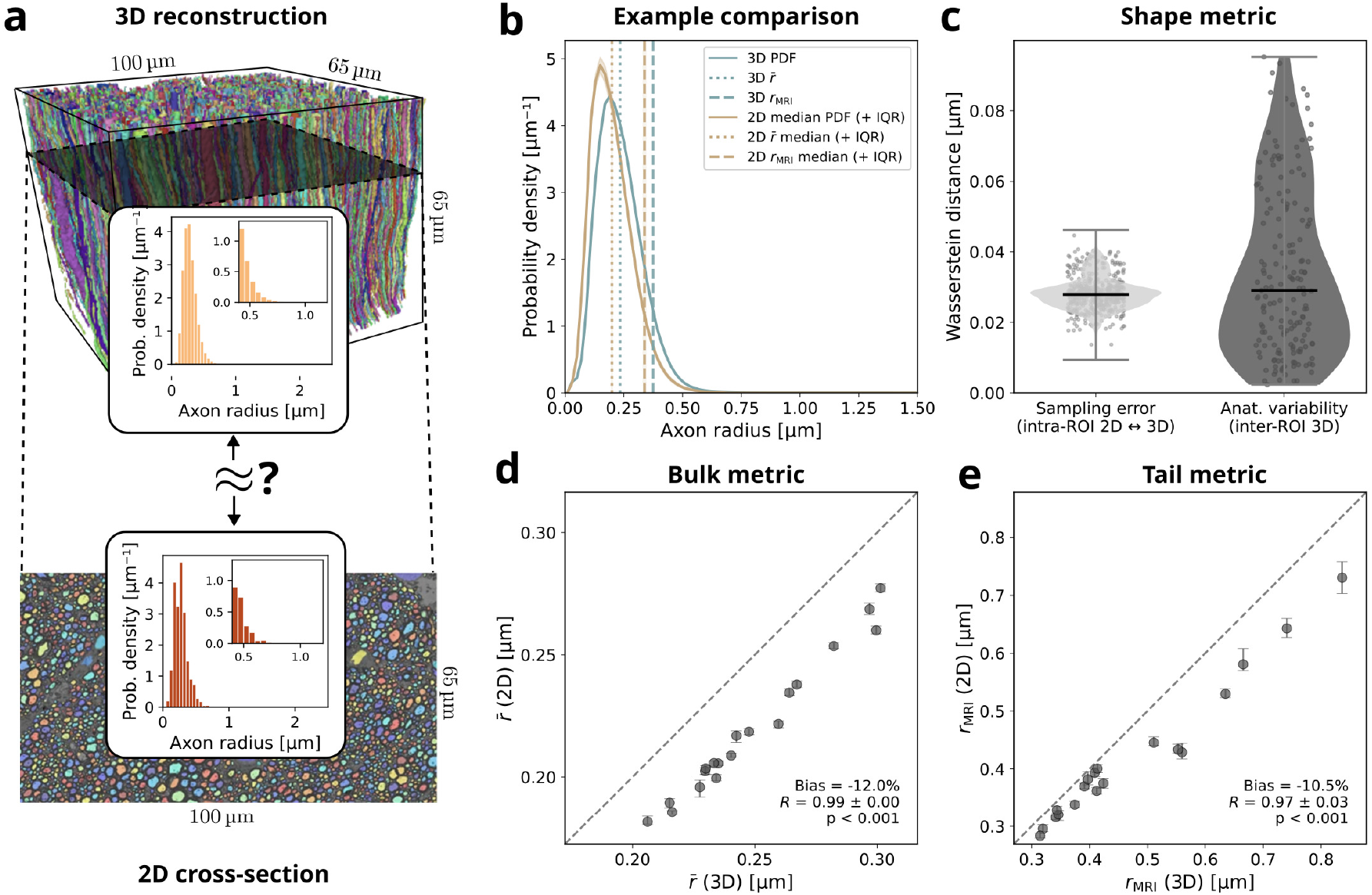
2D-derived axon radius distributions as 3D proxies. **(a)** 3D reconstruction of cingulum axons with corresponding 2D cross-section perpendicular to the bundle orientation, as typical for 2D histology. Insets show the corresponding axon radius distributions: 3D pools radii jointly across and along axons, whereas 2D captures only a single snapshot across axons. Do 2D distributions and derived ensemble metrics represent their 3D counterparts? **(b)** Example comparison for one ROI. Solid lines show 2D and 3D-based radius distributions; vertical lines mark ensemble average radii: the arithmetic mean 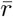 (dotted, see Eq. (9)) is sensitive to the distribution bulk; the MRI-visible axon radius *r*_MRI_ (dashed, see Eq. (10)) is sensitive to the distribution tail. For 2D, lines indicate median values across cross-sections, shaded bands represent IQR. **(c)** Distribution shape disagreement, quantified by Wasserstein distance (see Eq. (8)). The left distribution encodes all comparisons between distributions from 2D and their 3D counterpart, whereas the right distribution contextualizes this with inter-ROI distances between 3D distributions, reflecting the anatomical variation in the dataset. **(d–e)** Comparison of 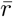 and *r*_MRI_ from 2D cross-sections versus 3D ground truth across all ROIs. Markers show median and IQR across 2D cross-sections for each ROI; dashed lines indicate identity. Pearson’s *R* and *p*-value were computed via Monte Carlo sampling.

Figure 3b compares the axon radius distribution from 2D cross-sections against the 3D counterpart for one representative ROI. The distributions are similar in shape, but the 2D distribution is shifted toward smaller radii, with a slightly larger bulk of small axons. To summarize the distributions, we computed two ensemble average radius metrics sensitive to different parts of the distribution: the arithmetic mean radius 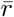 (see Eq. (9)), which reflects the distribution bulk, and *r*_MRI_ (see Eq. (10)), a radius metric that diffusion MRI is sensitive to, which is determined by the distribution tail. In correspondence with the shift between 2D and 3D distributions, both metrics are slightly lower in 2D than in 3D.

To quantify the 2D–3D correspondence across all ROIs, we assessed distribution shape via Wasserstein distance (see Eq. (8), Fig. 3c). The Wasserstein distance shows a consistent offset of approximately 0.04 µm, confirming the systematic bias of 2D cross-sections toward smaller radii. However, the spread between 2D and 3D-based distributions within each ROI is low, indicating that 2D-derived distributions are highly stable along the bundle. Moreover, this spread is much smaller than the Wasserstein distance differences due to anatomical variation across ROIs, suggesting that 2D cross-sections are sensitive to anatomical differences despite the bias.

This sensitivity to anatomical differences is confirmed by both ensemble average radii 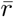 and *r*_MRI_(see Fig. 3d–e): 2D and 3D estimates are strongly correlated across ROIs (*R* ≥ 0.97, *p <* 0.001), despite 2D values being systematically lower (12 % for 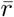, 10.5 % for *r*_MRI_).

The bias between distributions and metrics (see Fig. 3b–e) likely arises from the minor axis approximation in 2D versus perpendicular-to-centerline measurements in 3D, which can be mitigated by using circular equivalent radius, albeit at the cost of lower precision (see Fig. S2). Thus, the observed differences between 2D and 3D are driven by geometric approximations rather than a fundamental limitation of 2D sampling. Figure S3 underlines this notion, showing that 2D slice sampling is statistically equivalent to independent random sampling from the 3D distribution.

### Sample size requirements differ for bulk and tail metrics

Having established that 2D versus 3D sampling does not substantially change the observed axon radius distribution, sample size remains as an important consideration. To study sample size requirements across species, we conducted a subsampling analysis using massive axon radius distributions as population ground truth: the 3D rat data from previous sections (*>* 10^6^ radii per ROI, pooled across and along axons) and 2D human corpus callosum light microscopy (∼10^6^ axons per ROI).

Figure 4a contrasts sample sizes on a human corpus callosum light microscopy section: a whole sample (∼10^6^ axons) versus a subsample of 10^3^ axons, representative of typical human 2D microscopy datasets (e.g., Aboitiz et al. (1992); Liewald et al. (2014)). The resulting distributions agree in the bulk, but differ in the tail, where the subsample shows occasional spikes rather than the smooth decay with increasing radius of the whole sample.

**Figure 4.**
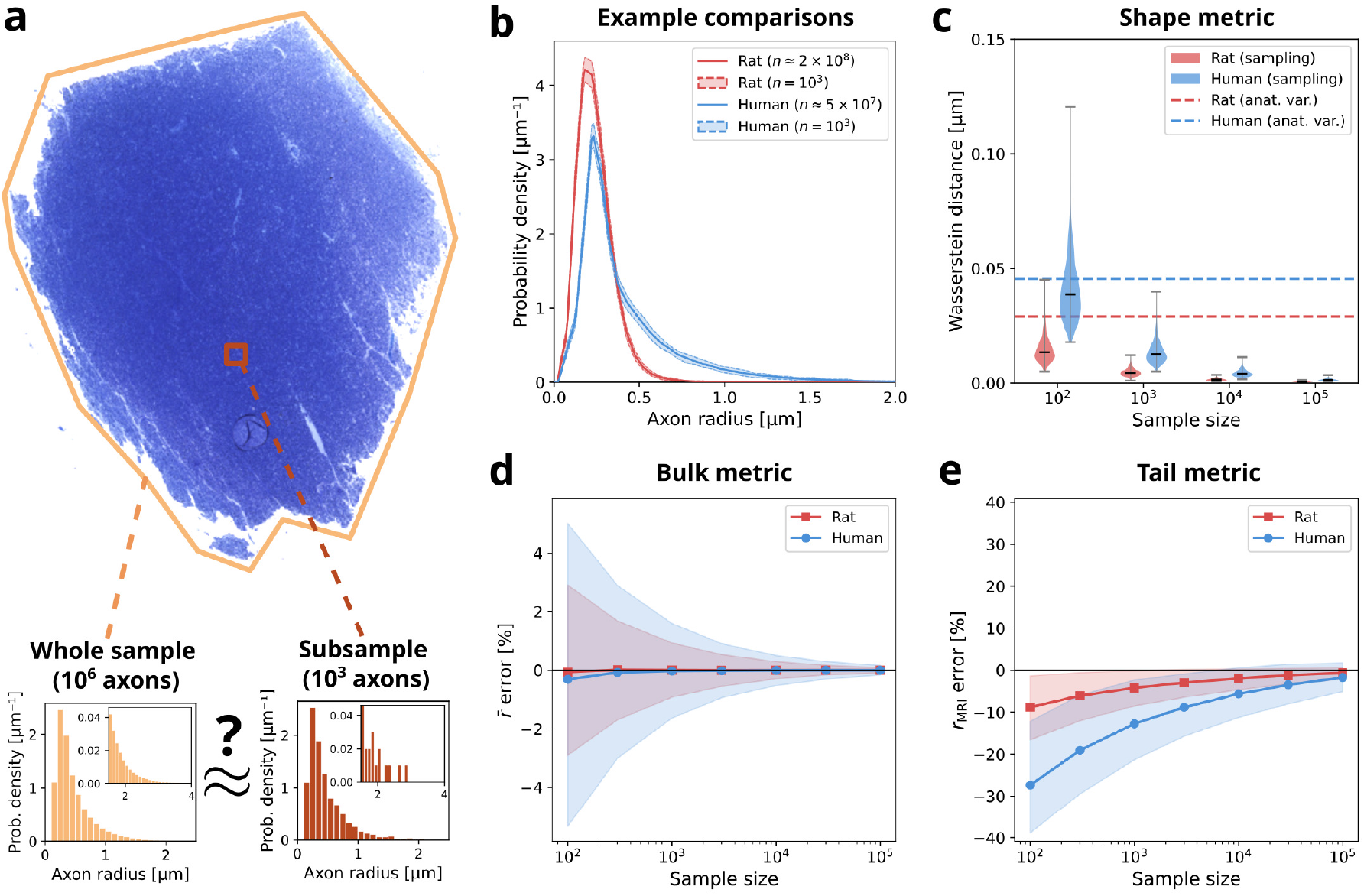
Sample size requirements for axon radius distributions. **(a)** Human corpus callosum light microscopy section illustrating whole sample (∼10^6^ axons) versus subsample (10^3^ axons). Histograms show probability density for both samples; insets emphasize the tail region. **(b)** Probability density pooled across all ROIs for human (blue; 35 ROIs, ∼5 × 10^7^ axons) and rat (red; 19 ROIs, ∼2 × 10^8^ radius samples). Solid lines show full data; shaded bands show IQR across subsamples at *n* = 10^3^. **(c)** Wasserstein distance (see Eq. (8)) as a function of sample size; vertical dashed lines indicate median inter-ROI anatomical variation for reference. **(d–e)** Relative error of ensemble average axon radii as a function of sample size: **(d)** arithmetic mean radius 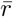 (see Eq. (9)); **(e)** MRI-visible axon radius *r*_MRI_ (see Eq. (10)). Markers show median; shaded regions indicate IQR (pooled across ROIs and subsamples). All simulations in (b–e) were conducted across 10^3^ subsamples per sample size.

Figure 4b shows the variation across repeated subsampling at sample size *n* = 10^3^ for both species. Variation is strongest for the large-axon tail in humans, reinforcing the impression from Fig. 4a that capturing the tail requires larger samples than the bulk.

To quantify these effects, Fig. 4c–e assess distribution shape and ensemble average radii as a function of sample size. The Wasserstein distance (see Fig. 4c) shows that at *n* = 10^2^, sampling errors can exceed the anatomical variation across ROIs (dashed lines), making it difficult to distinguish sampling noise from true anatomical differences. However, at *n* ≥ 10^3^, sampling error becomes much smaller.

For ensemble average radii (see Fig. 4d–e), 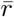 exhibits no bias and small variation (*<* 5 %) across all sample sizes for both species, suggesting that sample size error is below the systematic bias from minor axis approximation in 2D (|Bias| ≥ 10.5 %, see Fig. 3d-e). In contrast, *r*_MRI_ shows substantial underestimation at small samples: human data exhibits approximately 30 % bias at *n* = 10^2^, while rat data is less affected (∼ − 10 %). The *r*_MRI_ error converges toward zero for larger sample sizes, faster for rats (∼ − 10^3^ to 10^4^ axons) than for humans (10^5^ axons), underscoring that robust tail sampling requires adjustment to the species under study.

### Parametric distributions miss long tails

Beyond empirical distributions, parametric models offer a compact statistical representation of axon radius distributions. Given that previous attempts to determine the best parametric model (Sepehrband et al., 2016) used sample sizes (*n <* 10^4^) that may not capture long tails (see Fig. 4), we revisited parametric model identification at larger scale, using the 3D rat (pooled along and across axons) and 2D human data described above.

Fig. 5a-b show pooled axon radius distributions for rat and human, with candidate parametric fits overlaid. For rat, the candidate distributions cluster tightly and fit the data well. For human, the fits diverge and most distributions fail to represent both the mode and the tail.

**Figure 5.**
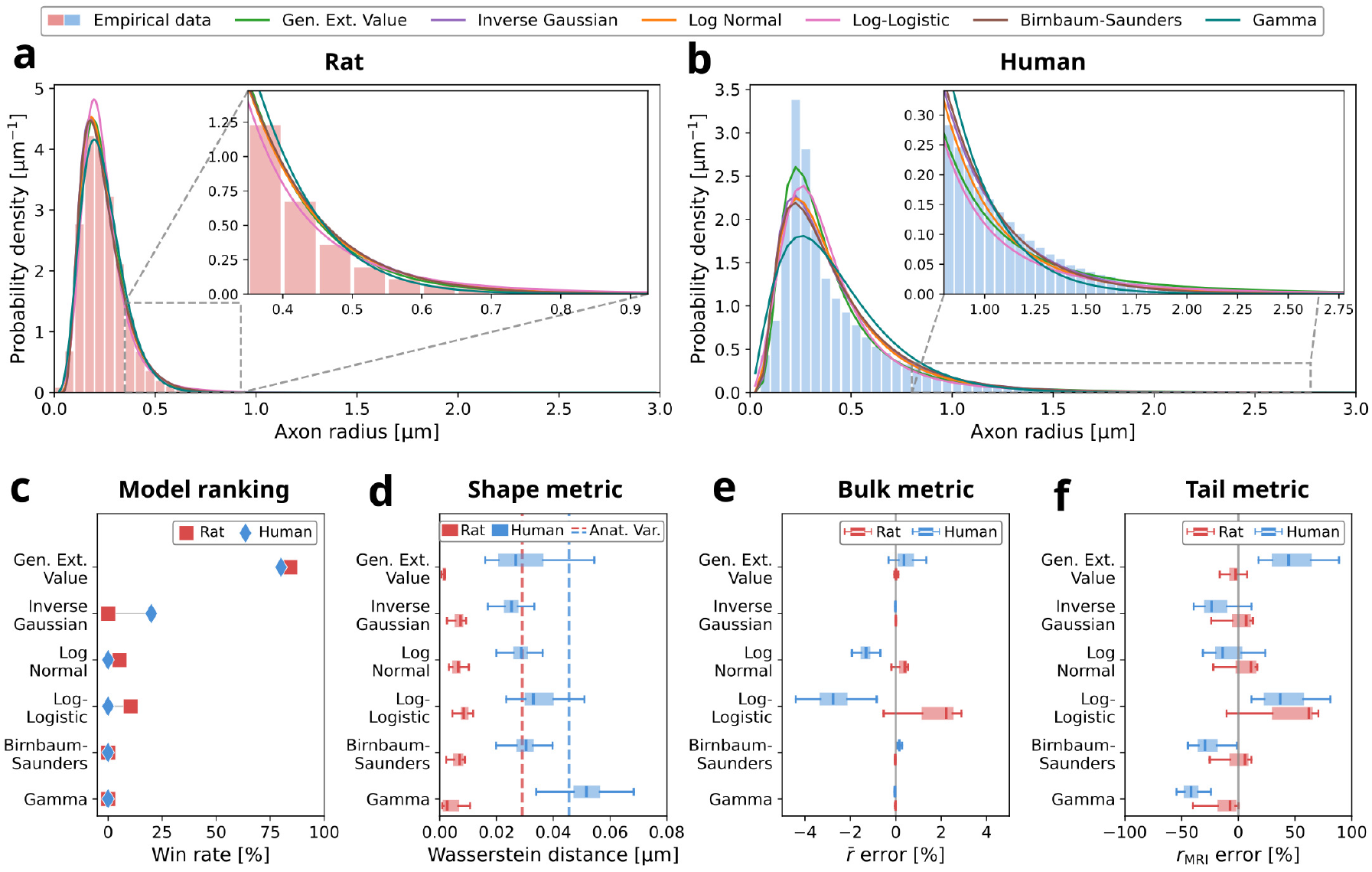
Parametric characterization of axon radius distributions. **(a–b)** Probability density pooled across ROIs for rat white matter and human corpus callosum with candidate parametric fits overlaid; insets emphasize the tail region. **(c)** Candidate model ranking. The win rate quantifies the proportion of ROIs where each candidate distribution model ranks best in terms of Akaike information criterion (AIC). **(d)** Wasserstein distance (see Eq. (8)) between fitted and empirical distributions; horizontal dashed lines indicate median inter-ROI anatomical variation for reference. **(e–f)** Relative error in ensemble average radii estimated from fitted distributions: **(e)** 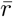; **(f)** *r*_MRI_. Box markers in (d–f) summarize variation across ROIs: median (vertical line), inter-quartile range (IQR, box), and 1.5× IQR (whiskers).

To rank candidate models for individual ROIs, we used the Akaike information criterion (AIC), which balances goodness-of-fit against model complexity. Fig. 5c shows the per-ROI win rate—the proportion of ROIs where each distribution achieves the lowest AIC. For both rat and human data, the generalized extreme value distribution wins in most ROIs, aligning with previous findings for long-tailed distributions in humans and monkeys (Sepehrband et al., 2016).

Fig. 5d assesses whether goodness-of-fit translates to accurate estimation of distribution shape (Wasserstein distance). Results echo earlier findings: distributions cluster closely for rat, while human preserves the model ranking from (c). Overall, shape mismatch between parametric fits and empirical data falls well below anatomical variation for rats (dashed line); for humans, values approach anatomical variation, especially for the Gamma distribution, indicating poorer fit.

The ensemble average radii 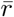 and *r*_MRI_ (see Fig. 5e-f), reveal the mechanism behind the shape mismatch. While 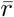, sensitive to the bulk of the distribution, shows low error across parametric distributions and species (*<* 5 %, Fig. 5e), *r*_MRI_, sensitive to the distribution tail, shows much larger errors (Fig. 5f). The effect is stronger in humans, where the heavier tail (see inset in Fig. 5b) leads to errors exceeding 25 % for most distributions. Notably, the largest error occurs for the generalized extreme value distribution in humans—the best-ranked model by AIC—suggesting that goodness-of-fit criteria may not predict tail accuracy.

## 3 Discussion

Using the largest available 3D and 2D white matter histology datasets, we examined along-axon radius variation and its functional implications, the robustness of axon radius distributions to histological sampling, and their parametric description. We found that, despite radius variation along individual axons, ensemble radius distributions remain stable along bundles, enabling 2D cross-sections to faithfully represent 3D bundle properties. Importantly, because along-axon variation has modest impact on conduction velocity, the representativeness of 2D cross-sections extends to conduction velocity predictions. In particular, large axons exhibit especially stable conduction, consistent with their key role in time-critical signaling. However, for diffusion MRI, along-axon variation has a larger impact, complicating interpretation. For histological sampling, we provide practical guidance: even small samples reasonably capture the distribution bulk, whereas larger samples are needed to characterize the long tail characteristic of human white matter. This long tail challenges parametric modeling of axon radius distributions, with standard parametric distributions failing to accurately capture tail-sensitive metrics.

### Axon bundles possess stable radius distributions

Much of the field’s knowledge about axon organization derives from 2D histology, including differences across tracts (Aboitiz et al., 1992; Liewald et al., 2014; Ruthig et al., 2025; Tomasi et al., 2012), species (Caminiti et al., 2009; Olivares et al., 2001; Wang et al., 2008) and health conditions (Sasaki and Maruyama, 1992; Wegiel et al., 2018). While the consistency of 2D-based findings, e.g., in across-species differences, lends credibility to the approach, we provide explicit validation that 2D radius distributions robustly describe the local 3D distribution of radii. Thereby, we provide strong empirical validation of previous hints from approximately 50 manually traced monkey axons (Andersson et al., 2020)—now across 19 fully segmented regions of interest (ROIs) totaling nearly 450,000 rat axons, including both traumatic brain injury (TBI) and control conditions.

The good capture of 3D axon radius distributions in 2D arises because radius variation is uncorrelated across axons and fluctuates only locally along axons, without systematic trends (Abdollahzadeh et al., 2025). Fig. S3 implicitly shows that these conditions are met by demonstrating that 2D cross-section sampling is statistically equivalent to random sampling from the pooled distribution across 2D cross-sections. The remaining systematic underestimation (∼12 %) arises from the minor axis approximation used in 2D, rather than from spatial structure within slices. The circular equivalent approximation could provide less biased estimates (see Fig. S2), but might require parametric filtering of obliquely cut axons for robust application.

### The cylinder model holds for conduction velocity

Conduction velocity predictions often assume cylindrical axons (Rushton, 1951; Schmidt and Knösche, 2019). Our results suggest this is a reasonable approximation: along-axon variation reduces conduction velocity by ∼4 % (median value) in the rat data investigated here, aligning with the small impact of focal axonal swellings in simulations (Kolaric et al., 2013). To put this in perspective: estimated conduction velocity in human corpus callosum exceeds superficial white matter by ∼20% (Ruthig et al., 2025). CoV-related reductions would not overshadow this, even for extreme CoV differences—ranging from ∼4% (CoV ≈ 0.27, the rat data investigated here) to ∼12% (CoV ≈ 0.5 (Tian et al., 2025)). Practically, the validity of both the cylinder model and 2D sampling reinforces findings from 2D histology-based conduction velocity research, ranging from tract-specific differences in conduction speed (Ruthig et al., 2025) to cross-species scaling where larger brains harbor giant axons that partially offset longer conduction distances (Caminiti et al., 2009).

### Large axons are optimized for stable conduction

We find that large axons exhibit less relative variation (CoV ≈ 0.15) than smaller axons (up to CoV ≈ 0.30). While the rat axons in this study are smaller than those in primates, limited 3D monkey corpus callosum data (50 axons, Andersson et al. (2020)) confirms low variation (CoV ≈ 0.12) for large axons with mean radius ≈1.5 µm, suggesting this stability generalizes across species and larger axons in organized tracts. This likely reflects the special constraints on large axons: functionally, they serve as “highways” for time-critical activity (Wang et al., 2008) and support high information rates that cannot be efficiently distributed across thinner axons (Perge et al., 2012); metabolically, their costs scale with diameter squared (Perge et al., 2009), favoring tighter optimization of these axons. This optimization of large axon conduction stability would follow a similar pattern established for g-ratio, where large axons cluster more tightly around the value that optimally balances conduction speed, energy efficiency, and volume constraints (Chomiak and Hu, 2009; Ruthig et al., 2025; Schmidt and Knösche, 2019; Tian et al., 2025).

### Diffusion MRI requires nuanced validation

Diffusion MRI offers a non-invasive window into axon microstructure, by measuring water diffusion within axons. In this modality the implications of along-axon radius variation are more nuanced than for conduction velocity.

One class of approaches targets the axon radius distribution—either attempting to estimate a scalar proxy (*r*_MRI_) (Burcaw et al., 2015; Veraart et al., 2020) or the full distribution (Assaf et al., 2008). These approaches rely on the cylinder model, assuming axons have a characteristic radius distribution. Under this model, the radius distribution and *r*_MRI_ are in principle well-specified according to our findings and can be validated with 2D histology. This is also supported by recent experimental correlation between in vivo MRI- and histology-based *r*_MRI_ in the human corpus callosum (Mordhorst et al., 2025a). That said, the diffusion MRI signal sensitive to axon radius coexists with diffusion MRI signal along axons in typical experimental regimes (Veraart et al., 2020). When falsely attributed to axon radius, this reduction in along-axon diffusion can reduce the specificity of radius estimates (Lee et al., 2020, 2024).

Yet, radius variation along axons is not merely a confounder but contains measurable anatomical information. A recent study has shown that along-axon morphology can directly be probed with diffusion MRI, enabling discrimination between TBI and control rats (Abdollahzadeh et al., 2025). Given that this approach explicitly models deviations from the cylinder model, 3D histology is needed for validation.

### Sample size requirements depend on application and species

While our findings suggest that 2D histological sampling of axon radius distributions is representative for 3D bundles, sample size remains a key consideration for designing histological studies. Our findings provide species- and application-dependent guidance for sample size selection, allowing researchers to balance sampling depth (within ROI) against width (e.g., across ROIs, donors, or conditions).

Capturing the bulk of the distribution (e.g., 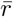) requires only modest sample sizes (∼10^3^ axons), typically met in most published human histology studies (e.g., Aboitiz et al. (1992); Barakovic et al. (2021); Caminiti et al. (2009); Graf von Keyserlingk and Schramm (1984); Liewald et al. (2014); Wegiel et al. (2018)). In contrast, capturing the tail is harder, requiring larger samples in humans (∼10^5^ axons) due to their long distribution tail. Such sample sizes are only reached in recent large-scale 2D (Mordhorst et al., 2022, 2025a; Ruthig et al., 2025) or 3D studies (Shapson-Coe et al., 2024; Tian et al., 2025).

For practical application, sample sizes with accurate bulk capture may suffice for ensemble conduction velocity comparisons, while tail capture is needed for robust estimates of maximum conduction velocities or *r*_MRI_ reference data.

### Mind the tail in parametric radius distribution modeling

Parametric distributions offer a compact statistical description of axon radius distributions for informing biophysical models—from compound action potential simulations to brain network dynamics and diffusion MRI (Assaf et al., 2008; Mitjans et al., 2023; Schmidt and Knösche, 2019; Schmidt and R. Knösche, 2022). Aligning with previous findings (Lee et al., 2019; Sepehrband et al., 2016), our data confirms that the generalized extreme value distribution is a leading candidate across species according to the Akaike Information Criterion (AIC), but we show that model selection does not ensure accurate representation. While rat data is reasonably captured according to all evaluated metrics, human data with its long distribution tail shows poorer representation. This tail seems particularly hard to represent, as illustrated by large errors in the tail-weighted *r*_MRI_—paradoxically highest for the AIC-selected generalized extreme value distribution. Also of note, the Gamma distribution, despite its popularity in conduction velocity (Oliveira et al., 2023) and diffusion MRI modeling (Assaf et al., 2008), also performs poorly. Overall, parametric modeling of axon radius distributions should be used with caution, particularly for applications that depend on accurate tail characterization.

### Limitations

Our 3D findings are derived from rat white matter, raising the question of whether they generalize to humans. However, monkey data—phylogenetically closer to humans—shows similar 2D–3D consistency and low along-axon variation for a limited dataset of 50 large axons (mean radius ≈ 1.5 µm, CoV ≈ 0.12, reported by Andersson et al. (2020)), suggesting these properties hold for species with longer distribution tails. This is further supported by recent correlation between in vivo MRI- and histology-based *r*_MRI_—a tail-weighted metric—in the human corpus callosum (Mordhorst et al., 2025a). We nonetheless suggest direct validation in large-scale human 3D axon reconstructions.

We pooled ROIs from sham-operated and TBI rats, treating each as an individual observation rather than comparing across conditions. However, the 2D–3D agreement remains close to the identity line across all ROIs, suggesting that the results are not driven by pathology-related differences.

We focused on along-axon radius variation but did not investigate axonal undulation (trajectory tortuosity). While Monte Carlo simulations suggest that caliber variation dominates over undulation for along-axon diffusion (Lee et al., 2020, 2024), undulation may also affect conduction velocity, though the gentle undulations reported in rat white matter (Abdollahzadeh et al., 2025) suggest a modest effect.

Our conclusions rest on observed radius distributions, which may differ from the true in vivo distributions. Tissue fixation causes shrinkage compared to in vivo conditions (Aboitiz et al., 1992; Dyrby et al., 2018; Tang et al., 1997; Yendiki et al., 2022), with recent work showing that shrinkage differs by axon radius (Skoven et al., 2025). Additionally, the human 2D light microscopy data cannot reliably resolve axons with radius below 0.3 µm (Mordhorst et al., 2022).

## 4 Conclusion

Traditionally, researchers have used 2D histological sections to study axon radius distributions, implicitly assuming cylindrical axons. While 3D histology has falsified this assumption, we show that ensemble-level conclusions—including radius distributions and conduction velocity predictions—from 2D remain valid. Beyond reinforcing decades of existing research, we provide guidance for efficient histological sampling across neuroscience applications.

## 5 Methods

### Human corpus callosum (2D light microscopy)

We analyzed 2D histological sections from the human corpus callosum, comprising 46 million axons across 35 ROIs from two tissue samples (one male, 61 years; one female, 60 years; postmortem delay 20–24 hours; see Fig. 1b, Mordhorst et al. (2025a)). Tailored for MRI validation, each ROI spans approximately 3 mm × 3 mm—comparable to in-vivo MRI voxels—yielding roughly one million axons per ROI. The tissue was immersion-fixed in 3% paraformaldehyde and 1% glutaraldehyde, and stained with toluidine blue. Myelinated axons were segmented from light microscopy images (0.11 µm/pixel resolution) using a deep-learning method (Mordhorst et al., 2022); axons with radius below ∼ 0.3 µm cannot be reliably detected at this resolution. Axon radii were estimated from the minor axis of ellipse fits to each cross-section; these radius measurements are provided in the public dataset (Mordhorst et al., 2025b). For full methodological details, see Mordhorst et al. (2022, 2025a).

### Rat white matter (3D electron microscopy)

We used publicly available 3D electron microscopy data (Abdollahzadeh et al., 2021; Sierra et al., 2021), comprising segmented myelinated axons from the rat corpus callosum and cingulum (see Fig. 1a). The dataset includes serial block-face scanning electron microscopy volumes from 5 adult male Sprague-Dawley rats: 3 with traumatic brain injury (TBI) and 2 sham-operated controls, imaged 5 months post-surgery. Both ipsilateral and contralateral hemispheres were sampled, yielding 10 volumes total. Each volume spans approximately 200 × 100 × 65 µm^3^ at 50 nm isotropic resolution, with approximately two-thirds corresponding to corpus callosum and one-third to cingulum, yielding 19 ROIs in total (10 volumes × 2 tracts, i.e., cingulum and corpus callosum^1^; see Fig. 1a). The dataset provides automated segmentations of myelinated axons obtained with the DeepACSON pipeline, which combines deep convolutional neural network-based semantic segmentation with cylindrical shape decomposition for instance segmentation (Abdollahzadeh et al., 2021). In total, approximately 450,000 myelinated axons are available.

#### ROI extraction

To separate each electron microscopy volume’s corpus callosum and cingulum populations (see Fig. 1a), we divided each volume into two spatial ROIs. For each ROI, we identified one of the image axes as the approximate bundle orientation, given that axons within each population are roughly aligned with coordinate axes.

#### Along-axon radius profiles

We extracted axon skeletons by tracing the medial axis via fast marching, using an implementation based on the original DeepACSON (Abdollahzadeh et al., 2021) implementation. We sampled the perpendicular cross-section every 0.05 µm along the skeleton and computed the radius of a circle with equivalent segmented area.

#### 2D cross-sections

To mimic traditional 2D histology, we extracted 2D slices along the image coordinate axis roughly aligned with the axon bundle orientation, accepting imperfect alignment as representative of experimental conditions. We estimated radii using the same minor-axis method as for the human data.

#### Outlier filtering

We manually inspected all axons with unusually large radii^2^ and rejected non-axonal regions such as segmentation artifacts or blood vessels.

### Along-axon radius variation and its impact

#### Quantification of variation

From the rat 3D radius profiles, we quantified variation along individual axons using the coefficient of variation:

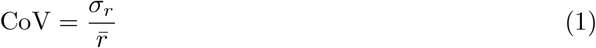

where *σ*_*r*_ is the standard deviation and 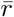 the mean radius along an axon’s trajectory. To ensure robust CoV estimates, we restricted analysis to axons exceeding 20 µm arc length, providing sufficient sampling along each axon’s trajectory.

#### Conduction velocity reduction

The conduction velocity of myelinated axons depends on the local radius *r* and g-ratio *g* as (Rushton, 1951; Schmidt and Knösche, 2019)

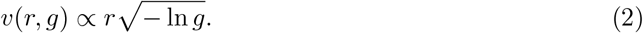

For an idealized cylindrical axon with 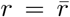 along the axon trajectory, this gives a uniform conduction velocity *v*_ideal_. For a realistic axon, however, variation of *r* and *g* along the axon trajectory leads to non-uniform velocity. The effective velocity over a segment of length *L* is

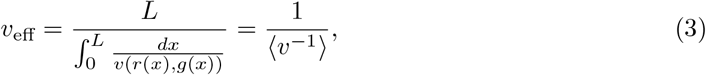

and the conduction velocity reduction relative to the ideal cylinder is

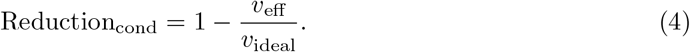

We evaluated Eq. (4) from the sampled radius profiles along axon trajectories. To this end, we assumed constant myelin thickness *d*_*m*_ along the axon (Andersson et al., 2020; Behanova et al., 2022) and a mean g-ratio of 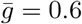 near the conduction-optimal value (Chomiak and Hu, 2009; Rushton, 1951), so that the local g-ratio depends only on the local radius:

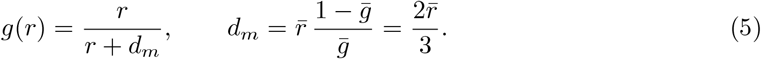

Under these assumptions, Eq. (4) can be expressed in terms of CoV via Taylor expansion (see Appendix S1):

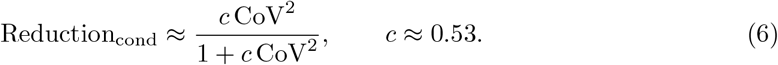

#### Along-axon diffusion reduction

For along-axon diffusion, caliber variations reduce the effective diffusivity compared to an ideal cylinder as (Lee et al., 2020):

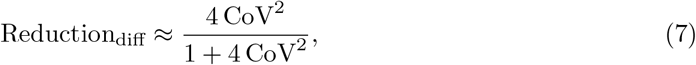

The factor of 4 relative to conduction velocity arises because diffusion is governed by crosssectional area (*A* ∝ *r*^2^), so CoV^2^(*A*) = 4 CoV^2^(*r*).

### Axon radius distribution analyses

#### 2D-derived distributions as 3D proxies

To compare 2D and 3D representations of axon radius distributions, we extracted multiple 2D cross-sections perpendicular to the bundle from each 3D ROI. For each cross-section, we computed a 2D radius distribution (*P*_2D_) and compared it to the ROI’s 3D reference distribution (*P*_3D_), which pools radii measured along axon trajectories across all axons within the ROI. We quantified agreement using complementary metrics below.

To assess distribution shape, we used the Wasserstein distance (specifically the 1-Wasserstein or earth mover’s distance, Kantorovich (1960)):

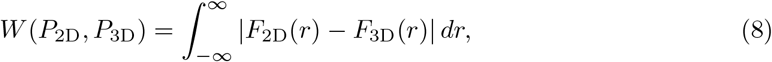

where *F*_2D_ and *F*_3D_ are the cumulative distribution functions corresponding to *P*_2D_ and *P*_3D_. As a reference for interpreting 2D–3D deviations, we computed the anatomical variation—defined as the median pairwise Wasserstein distance between 3D distributions from different ROIs.

To assess ensemble average radii, we compared two metrics: the arithmetic mean radius

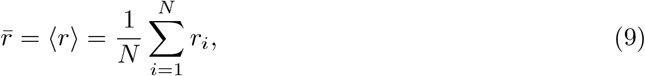

which provides a bulk-weighted measure of typical axon size, and the MRI-visible axon radius (Burcaw et al., 2015; Veraart et al., 2020)

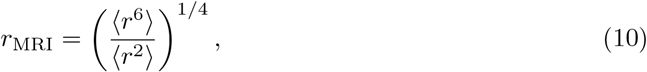

which weights larger axons more heavily and is particularly sensitive to the distribution tail.

To quantify systematic offset, we computed the normalized mean bias error

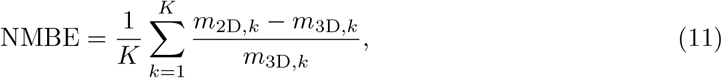

where *m* denotes the metric of interest (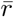 or *r*_MRI_), *k* indexes ROIs, and *K* is the number of ROIs.

To assess correlation between 2D and 3D estimates of 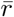 and *r*_MRI_, we mimicked a typical 2D histological study by sampling a single *P*_2D_ per ROI and computing Pearson’s *R* with the corresponding *P*_3D_ values across ROIs. We repeated this over 10,000 iterations, with *p*-values derived from Monte Carlo permutation testing, estimating *p* as the fraction of permutations where the permuted *R* exceeded the observed *R*.

#### Sample size requirements

To assess how sample size affects estimation accuracy, we performed a subsampling analysis using the 3D radius distributions from rat and 2D human radius distributions as population ground truth (both ∼ 10^6^ or more radius samples per ROI). From each ROI’s full sample, we drew random subsamples of sizes *n* ∈ [10^2^, 10^5^] with 1,000 repetitions per *n*. For each subsample, we quantified deviation from the full sample using *W* for distribution shape, and relative error in 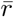 and *r*_MRI_. This random subsampling assumes spatially uncorrelated axon radii, which is supported by our finding that 2D slice sampling is statistically equivalent to random sampling from the pooled distribution (see Fig. S3).

#### Parametric characterization

We evaluated six candidate parametric distributions, identified as top candidates by Sepehrband et al. (2016): gamma, lognormal, generalized extreme value, inverse Gaussian, Birnbaum-Saunders, and log-logistic. For each distribution, we estimated parameters per ROI using binned maximum likelihood and evaluated goodness-of-fit using the Akaike Information Criterion (AIC). We quantified fit quality using *W* between fitted and empirical distributions, and bias in 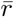 and *r*_MRI_ by comparing values obtained via numerical integration of the fitted PDF, evaluated from 0 µm to the 99.999th percentile of pooled radii per species, to empirical values.

## Data availability

The 3D electron microscopy rat data and its segmentation is publicly available at https://etsin.fairdata.fi/dataset/f8ccc23a-1f1a-4c98-86b7-b63652a809c3 (Sierra et al., 2021). The axon radius distributions from 2D human light microscopy data are publicly available at https://zenodo.org/records/17431227 (Mordhorst et al., 2025b).

## Code availability

All code to reproduce the results and figures from the datasets above is publicly available at https://github.com/quantitative-mri-and-in-vivo-histology/axon_caliber_variation.

## Author Contributions

L.M.: conceptualization, methodology, software, data curation, investigation, formal analysis, visualization, writing of the original draft. N.W.: funding acquisition, review and editing. M.M.: funding acquisition, review and editing. S.M.: supervision, funding acquisition, review and editing.

## Acknowledgement

We thank Tilo Reinert for insightful discussions.

## Supplementary Material

### S1 Derivation of the conduction velocity reduction formula

Here, we derive Eq. (6) from the main text.

#### Velocity under constant myelin thickness

Substituting the g-ratio model from Eq. (5) into the velocity relation Eq. (2) gives

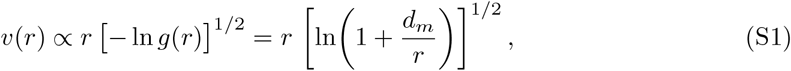

where the velocity now depends only on the local radius *r* (since *d*_*m*_ is constant along the axon (Andersson et al., 2020; Behanova et al., 2022)).

#### Reduction in terms of slowness

Defining the slowness *h*(*r*) = 1*/v*(*r*) (time per unit length), the transit time over a segment of length *L* is *t* = *L* ⟨*h*(*r*)⟩, so that *v*_eff_ = *L/t* = 1*/* ⟨*h*(*r*)⟩.

The reduction from Eq. (4) then becomes

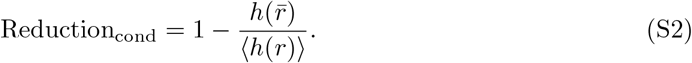

#### Taylor expansion

Writing 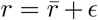 with ⟨*ϵ*⟩ = 0 and 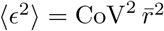, a second-order Taylor expansion of *h* around 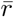 gives

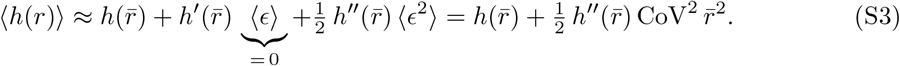

Substituting into Eq. (S2) gives

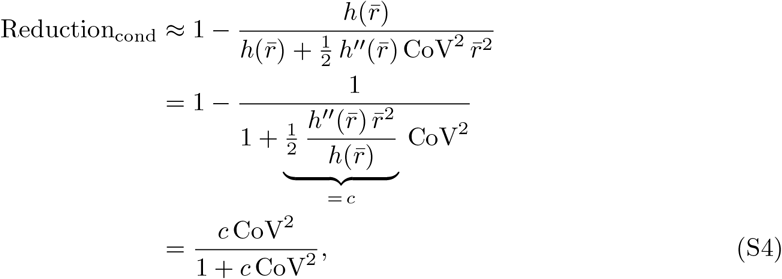

where *c* depends only on the dimensionless ratio 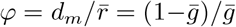, not on 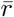 itself. Computing 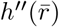 from Eq. (S1) gives the closed-form expression

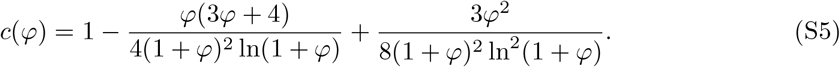

For 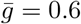 (Chomiak and Hu, 2009; Rushton, 1951) (i.e., *φ* = 2*/*3), this yields *c* ≈ 0.53.

### S2 Absolute radius variation saturates for thick axons

**Figure S1:**
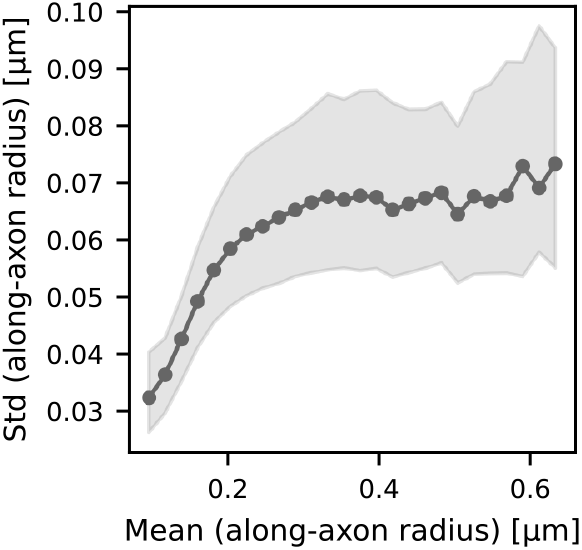
Absolute radius variation saturates for thick axons. Standard deviation of radius along individual axons as a function of along-axon mean radius. Axons are binned by along-axon mean radius; line shows median, shaded region shows IQR. To robustly quantify along-axon radius variation, we computed statistics for axons exceeding 20 µm arc length (*n* ≈ 150,000); bins were constrained to a minimum of 50 axons. The standard deviation saturates for thick axons, explaining why the coefficient of variation (CoV) in Fig. 2d decreases for thick axons.

### S3 Circular equivalent radius reduces bias but degrades precision

**Figure S2:**
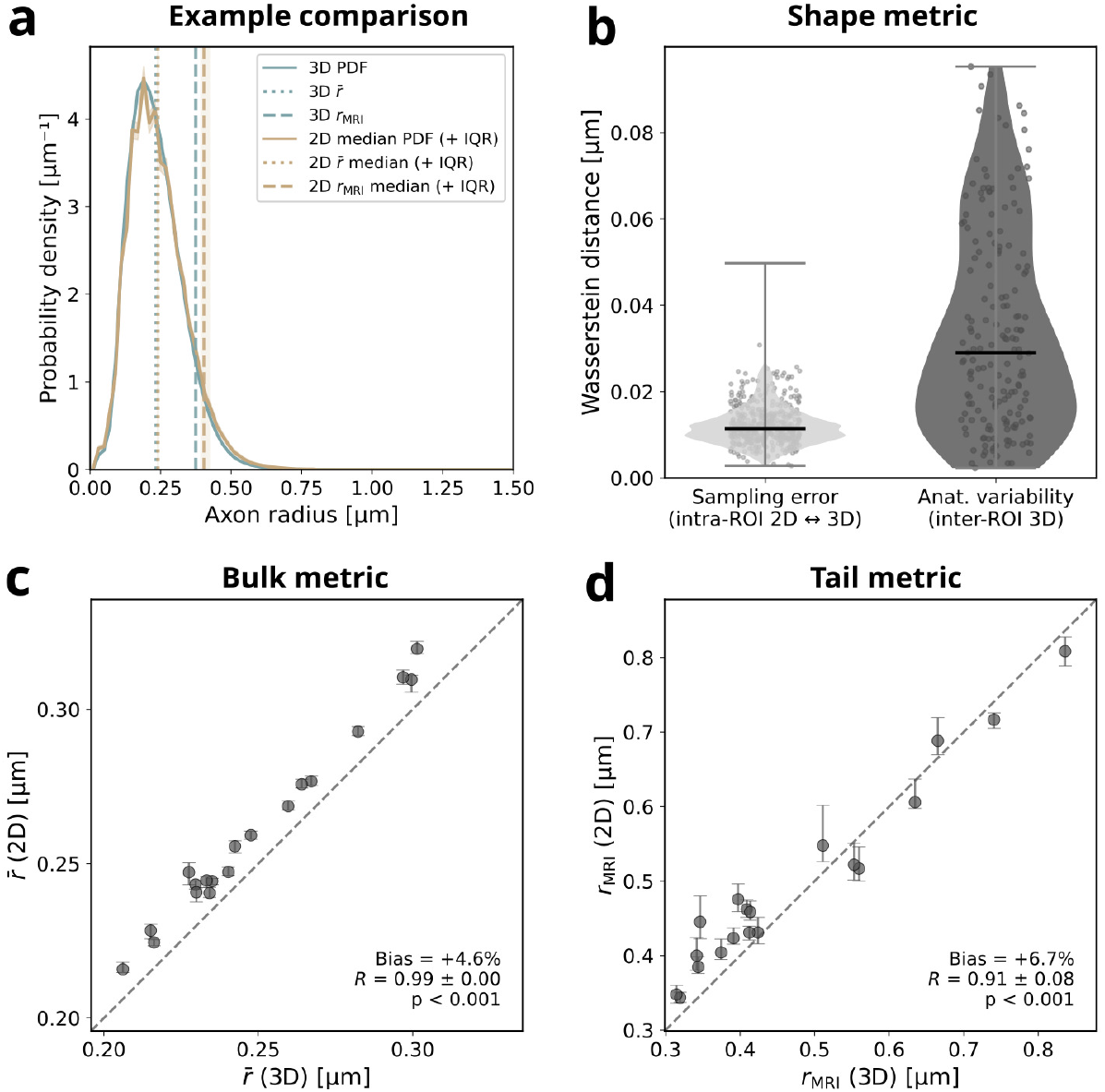
Circular equivalent radius eliminates bias but reduces precision. Same analysis as Fig. 3b–e, but using circular equivalent radius in 2D instead of the minor axis approximation. **(a)** Example comparison for one ROI. Solid lines show 2D and 3D-based radius distributions; vertical lines mark ensemble average radii: the arithmetic mean 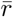 (dotted, see Eq. (9)) is sensitive to the distribution bulk; the MRI-visible axon radius *r*_MRI_ (dashed, see Eq. (10)) is sensitive to the distribution tail. For 2D, lines indicate median values across cross-sections, shaded bands represent IQR. **(b)** Distribution shape disagreement, quantified by Wasserstein distance (see Eq. (8)). The left distribution encodes all comparisons between distributions from 2D and their 3D counterpart, whereas the right distribution contextualizes this with inter-ROI distances between 3D distributions, reflecting the anatomical variation in the dataset. **(c–d)** Comparison of 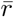 and *r*_MRI_ from 2D cross-sections versus 3D ground truth across all ROIs. Markers show median and IQR across 2D cross-sections for each ROI; dashed lines indicate identity. Pearson’s *R* and *p*-value were computed via Monte Carlo sampling.

### S4 2D slice-wise sampling is statistically equivalent to random sampling

**Figure S3:**
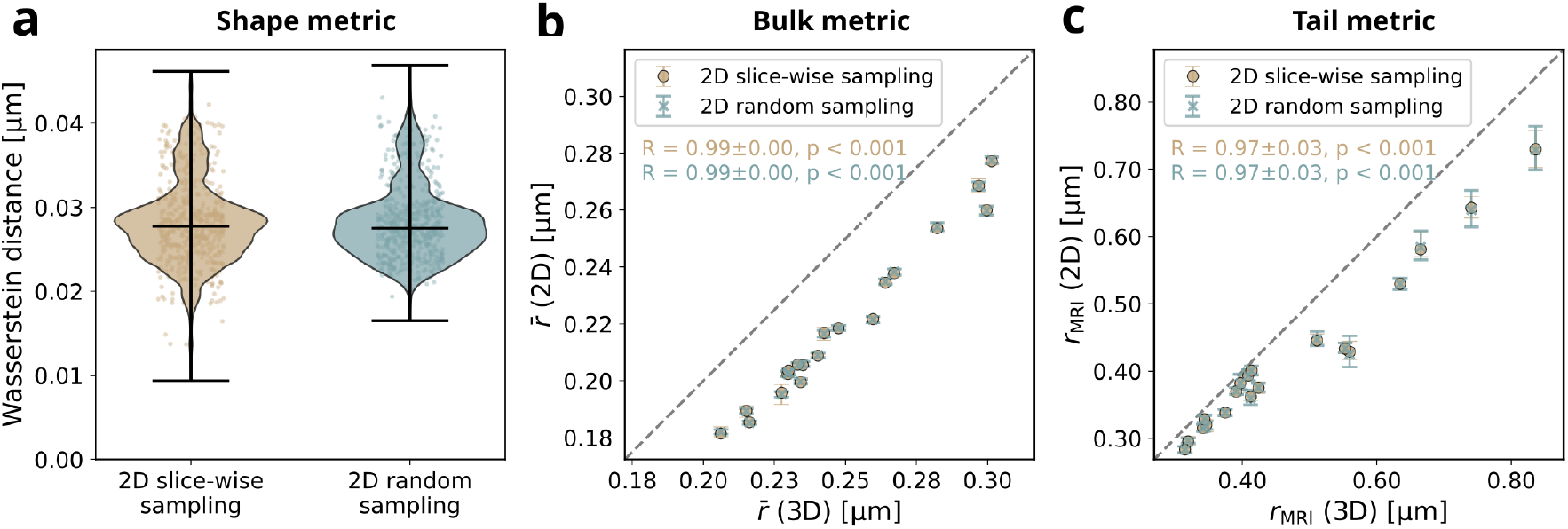
2D slice-wise sampling is statistically equivalent to random sampling. Monte Carlo comparison (10,000 iterations) of actual 2D slices versus random sampling without replacement from 2D radii pooled across the whole ROI. Each random sample was matched in sample size with slice-wise samples. **(a)** Wasserstein distance between sampled distribution and joint 3D ground truth. Violin plots show distribution across all ROI-iteration pairs. **(b)** Mean radius 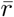 from 2D samples versus 3D reference. Points show median per ROI; compared to Fig. 3d–e, error bars here show IQR across Monte Carlo iterations, not across all 2D cross-sections of one ROI. **(c)** MRI-visible radius *r*_MRI_, format as in (b). The remarkable similarity between slice-wise and random sampling demonstrates that 2D-to-3D deviations do not arise from spatial structure within slices but rather from geometric approximations used in 2D histological sampling (see Fig. S2).

## Footnotes

1 corpus callosum data was not available for one volume, yielding 19 instead of 20 ROIs

2 Axons with *r*_MRI_ ≥ 1.5 µm (see Eq. (10))

